## Supplementary material for "CHC22 clathrin functions in the early secretory pathway by two-site interaction with SNX5 and p115": Fig. EV5: EV5(R2).docx

Table S5 - Buffers and Reagents used for in vitro binding, competition assays and immunoprecipitations.

| Buffer Buffer Composition | |
| --- | --- |
| IP lysis buffer | 20 mM HEPES pH 7.5, 150 mM NaCl, 1 mM EDTA, 1 mM EGTA,  10% vol/vol Glycero1, and 0.25% vol/vol Triton X-100 |
| Fractionation buffer | 20 mM HEPES pH 7.5, 400 mM sucrose, 1 mM EDTA, 1 mM PMSF, 1 mM Na_3_VO_4_ |
| TD binding buffer | 50 mM Tris pH 8, 300 mM NaCl, 5% glycerol, 25 mM imidazole |
| TD lysis buffer | 50 mM Tris pH 8, 300 mM NaCl, 5% glycerol, 25 mM imidazole, 1 mM PMSF, EDTA-free protease inhibitor |
| TD wash buffer | 50 mM Tris pH 8, 300 mM NaCl, 5% glycerol, 50 mM imidazole |
| TD elution buffer | 50 mM Tris pH 8, 300 mM NaCl, 5% glycerol, 250 mM imidazole |
| TD size exclusion buffer | 50 mM Tris pH 8, 150 mM NaCl, 5% glycerol |
| TxD binding buffer | 20 mM HEPES pH 7.5, 500mM NaCl, 20mM imidazole |
| TxD lysis buffer | 20 mM HEPES pH 7.5, 500mM NaCl, 20mM imidazole, 1 mM PMSF,  EDTA-free protease inhibitor tablet (Roche) |
| TxD wash buffer | 20 mM HEPES pH 7.5, 500mM NaCl, 50mM imidazole |
| TxD elution buffer | 20 mM HEPES pH 7.5, 500mM NaCl, 500mM imidazole |
| TxD size exclusion buffer | 20 mM HEPES pH 7.5, 200mM NaCl, 3 mM BME |
| SNX binding buffer | 20 mM Tris pH 8, 400 mM NaCl, 10% glycerol, 2 mM EDTA, 1 mM  BME |
| SNX lysis buffer | 20 mM Tris pH 8, 400 mM NaCl, 10% glycerol, 2 mM EDTA, 1 mM  BME, 1 mM PMSF, EDTA-free protease inhibitor |
| SNX5 elution buffer | 20 mM Tris pH 8, 400 mM NaCl, 10% glycerol, 2 mM EDTA, 1mM  BME, 10 mM reduced glutathione |
| SNX1 wash buffer | 20 mM Tris pH 8, 400 mM NaCl, 10% glycerol, 2 mM EDTA, 1 mM  BME, 10 mM imidazole |
| SNX1 elution buffer | 20 mM Tris pH 8, 400 mM NaCl, 10% glycerol, 2 mM EDTA, 1 mM  BME, 100 mM imidazole |
| SNX size exclusion buffer | 20mM Tris pH 8, 200mM NaCl, 10% glycerol, 2 mM EDTA, 1mM  BME |
| His purification buffer | 50 mM Tris pH 8, 500 mM NaCl, 10 mM imidazole, 5% glycerol, 1 mM PMSF, EDTA-free protease inhibitor |
| His PD buffer | 50 mM Tris pH 8, 300 mM NaCl, 2 mM EDTA, 1 mM BME, 2% BSA |
| His PD wash buffer | 50 mM Tris pH 8, 300 mM NaCl, 2 mM EDTA, 1 mM BME |
| SNX5 blocking buffer | 20 mM Tris pH 8, 400 mM NaCl, 10% glycerol, 2 mM EDTA, 1 mM  BME, 5% BSA, 1 mM PMSF |
| SNX1 PD buffer | 20 mM Tris PH8, 200 mM NaCl, 10% glycerol, 2 mM EDTA, 1 mM  BME, 5% BSA, 1 mM PMSF |
| TxD PD buffer | 20 mM HEPES pH 7.5, 300 mM NaCl, 3 mM BME, 5% BSA, 1mM PMSF |
| TxD PD wash  buffer | 20 mM HEPES pH 7.5, 300 mM NaCl, 3 mM BME, 1mM PMSF |
| GST adaptor buffer | 20 mM HEPES pH 7.5, 150 mM NaCl, 2 mM BME, 1 mM EDTA, 1 mM PMSF, EDTA-free protease inhibitor |
| CHC22 PD buffer | 20 mM HEPES pH 7.5, 300 mM NaCl. 2mM BME, 1mM EDTA, 50 mM NaSCN, 0.05% NP-40, 5% BSA |
| CHC22 PD wash  buffer | 20 mM HEPES pH 7.5, 300 mM NaCl. 2mM BME, 1mM EDTA, 50 mM NaSCN, 0.05% NP-40 |
| CHC17 blocking buffer | 20 mM HEPES pH 7.5, 150 mM NaCl, 0.05% NP-40, 2% ovalbumin |
| CHC17 PD buffer | 20 mM HEPES pH 7.5, 150 mM NaCl, 0.05% NP-40 |
| GFP IP lysis buffer | 10 mM Tris pH 7.5, 150 mM NaCl, 0.5 mM EDTA, 0.5% NP-40, 10% glycerol, EDTA-free protease inhibitor, 1 mM PMSF and 4 mM Na_3_VO_4_ |
| GFP IP dilution buffer | 10 mM Tris pH 7.5, 150 mM NaCl, 0.5 mM EDTA, 10% glycerol,  EDTA-free protease inhibitor, 1 mM PMSF and 4 mM Na_3_VO_4_ |
