## Supplementary figures and images for "CHC22 clathrin functions in the early secretory pathway by two-site interaction with SNX5 and p115"

### Fig. EV1

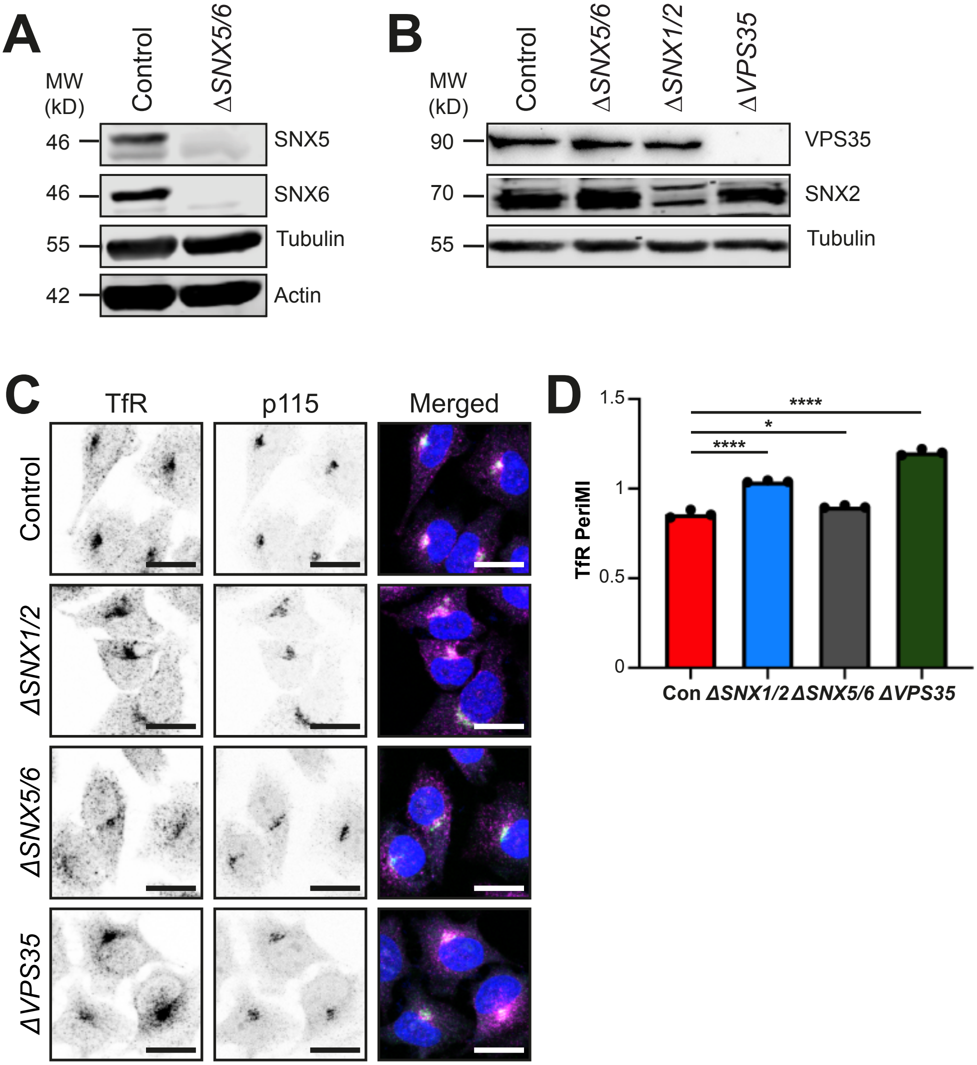

### Fig. EV2

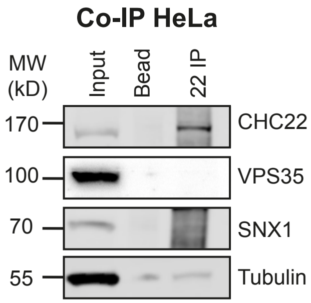

### Fig. EV3

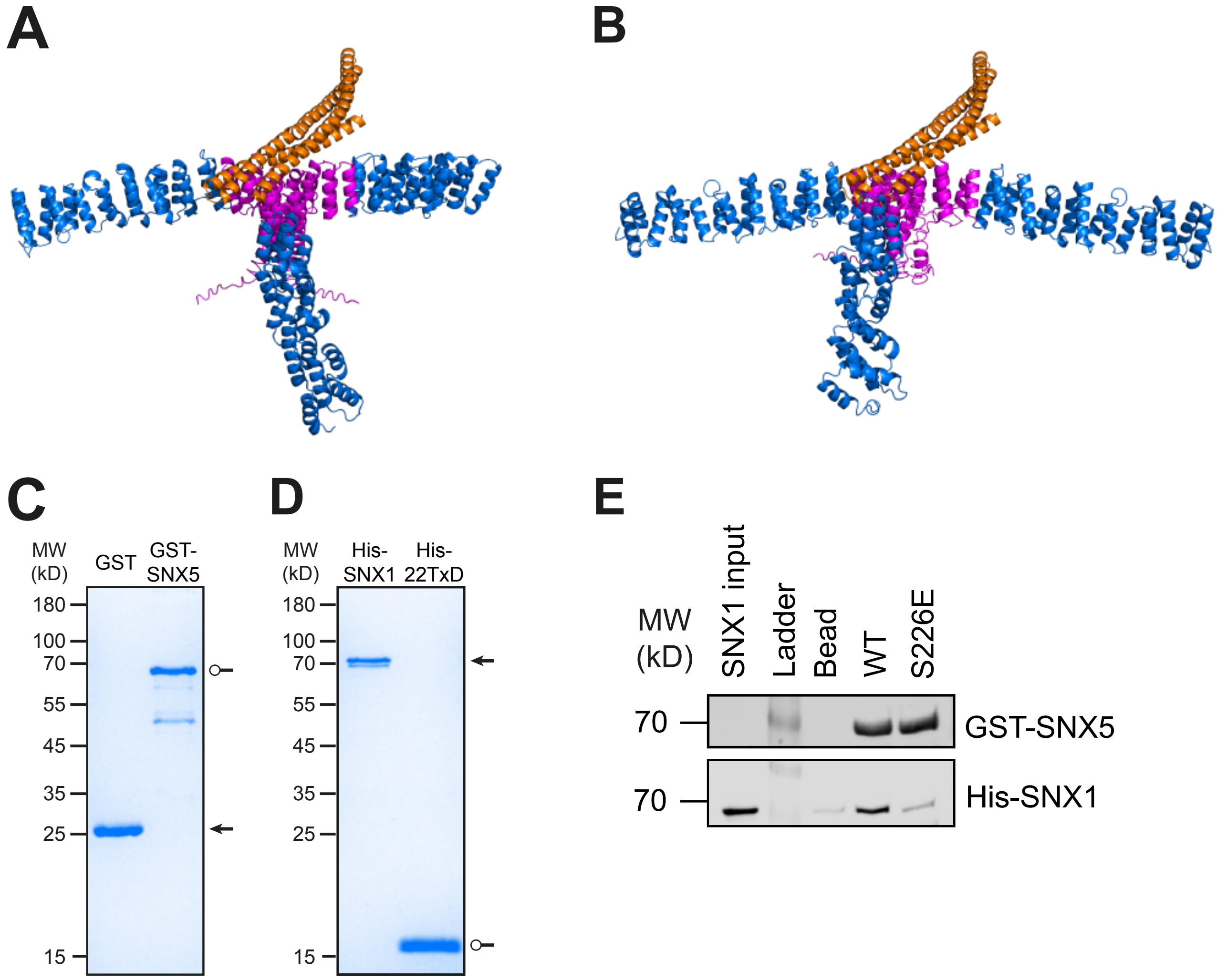

### Fig. EV4

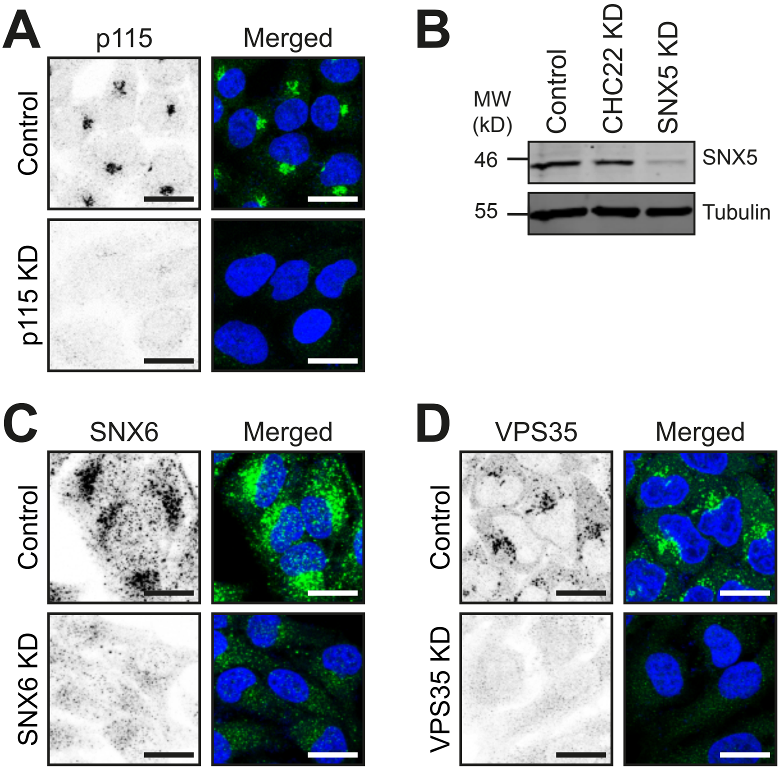
